# Spinal dI3 neurons are involved in sustained motor adaptation elicited by low-threshold cutaneous afferents

**DOI:** 10.1101/2025.09.22.677841

**Authors:** Emam U. Khan, Shahriar Nasiri, Sarah A. Chiasson, Lauren J. Couvrette, Alex M. Laliberte, Tuan V. Bui

## Abstract

Adaptation of muscle activity to meet a certain target or intention is traditionally attributed to supraspinal structures. However, evidence is mounting that this process can occur within the spinal cord through intrinsic plasticity and circuit reorganization. Here, we investigate the role of a class of excitatory spinal interneurons, called dorsal interneuron 3 (dI3), in the acquisition of novel motor behaviors independent of supraspinal input. Using a real-time closed-loop stimulation paradigm in spinalized mice, we promoted a persistent adaptation in the hindlimb position to be higher than its resting level by delivering saphenous nerve stimulation contingent on toe elevation. The stimulation intensities were calibrated to selectively recruit low-threshold mechanoreceptors (LTMRs). To test the contribution of dI3s in this motor adaptation, inhibitory DREADD (hM4Di) receptors were expressed in Isl1⁺/Vglut2⁺ cells, achieving reversible, cell-type-specific silencing of dI3s. Our results demonstrate that stimulation of cutaneous inputs to the spinal cord contingent on a certain positional goal can generate sustained changes in motor activity, in this case, in the form of elevation of toe position above a preset vertical threshold. Chemogenetic silencing of dI3s abolished this motor adaptation induced by activation of LTMRs. These findings indicate that dI3 activity is essential for a particular type of motor adaptation driven primarily by LTMR input.

**NEW & NOTEWORTHY:** We developed a real-time, closed-loop stimulation paradigm in spinalized mice using kinematic video tracking to trigger electrical stimulation of the saphenous nerve. We discovered that low-threshold stimulations targeting non-nociceptive cutaneous afferents can elicit sustained motor adaptations independently from supraspinal input. Furthermore, using two chemogenetic techniques to transiently inhibit a population of spinal neurons, called dI3s, we found that these neurons are crucial for integrating these low-threshold stimuli to elicit sustained changes in motor behaviour.

## INTRODUCTION

The role of spinal circuits in mediating rapid adaptation of on-going movements to perturbations is well-established. Spinal reflexes are classic examples of such rapid adaptations. More sustained adaptation of motor activity has long been thought of being the purview of supraspinal regions such as the cerebellum (Bastian et al., 1996; Smith and Shadmehr, 2005; Morton and Bastian, 2006; Synofzik et al., 2008; Taylor et al., 2010; Nguyen-Vu et al., 2013). However, a handful of studies suggest that with repetitive sensory stimulation, the spinal cord is capable of generating sustained motor adaptation (Carp et al., 2001; Carp & Wolpaw, 1994, 1995; Ferguson et al., 2012; Wolpaw, 2007; Wolpaw & Lee, 1989; Zhong et al., 2012).

Buerger and Fennessy (1970, 1971) developed a limb withdrawal paradigm in spinalized rats to demonstrate that this sustained adaptation is facilitated solely by the spinal cord. In this paradigm, an electric stimulus was applied to the hindlimb every time its position reached below a set threshold. Repetitions of this stimulation contingent on the position of the limb led to the maintenance of a flexed position above the threshold (Buerger and Fennessy, 1970, 1971). Conversely, control mice which received the same stimulations that were not contingent on their limb position, were unable to maintain this elevated limb position (Buerger and Fennessy, 1970, 1971). Subsequently, it was found that these sustained motor adaptations are mediated by synaptic plasticity through activation of NMDA receptors, BDNF signalling, and protein synthesis (Crown et al., 2002; Joynes et al., 2004; Baumbauer et al., 2009); however, the neuronal populations and circuitry critical for these behaviours were largely unknown until recently.

The development of genetic tools have allowed us to characterize and manipulate the activity of genetically distinct neuronal populations in the spinal cord (Zhang et al., 2008, 2014; Bui et al., 2013; Dougherty et al., 2013; Delile et al., 2019; Russ et al., 2021), with diverse connectivity and function. Using these approaches, Lavaud et al. (2024) investigated the role of several neuronal populations in a similar motor adaptation paradigm. They utilized a closed-loop muscle electrical stimulation protocol to train spinalized mice to maintain their toe above a threshold height. Afterwards, the importance of specific spinal neuron populations was evaluated by ablation of several classes of molecularly-defined spinal neurons (Lavaud et al., 2024).

Notably, ablation of dI4 neurons, compromised adaptation of toe height (Lavaud et al., 2024). These neurons are comprised of multiple subpopulations of inhibitory neurons, some of which are involved in the gating of nociceptive sensory feedback through presynaptic inhibition (Duan et al., 2014; Koch et al., 2017; Zimmerman et al., 2019). Lavaud et al. (2024) found that dI4s presynaptically inhibit nociceptive input (A-δ and C fibers) onto nociceptive-selective neurons and did not presynaptically inhibit neurons integrating both proprioceptive and nociceptive inputs. These nociceptive-selective neurons are located in the superficial dorsal horn lamina I and II. In addition to their central role in nociception (Agashkov et al., 2019), these superficial laminae neurons are also involved in motor behaviours such as scratching or withdrawal (Gatto et al., 2021) whereas neurons in lamina III-V integrating both proprioceptive and cutaneous and high-threshold sensory feedback, called wide dynamic range neurons (Woller et al., 2017), are involved in corrective behaviours (Gatto et al., 2021).

Lavaud et al. (2024) also putatively identified the downstream neurons integrating these multiple modalities of sensory input through ablation of Tlx3 expressing neurons. These Tlx3-positive neurons, comprised of premotor dI3 and dI5 neurons, have been shown to be involved in high threshold mechanoreceptive and nociceptive processing pathways (Goetz et al., 2015; Xu et al., 2008), and their ablation was found to impair the acquisition of this motor adaptive behaviour (Lavaud et al., 2024). Lastly, it was found that an inhibitory populations of ventral interneurons, called Renshaw cells, facilitate the recall of this learned motor behaviour by modulating the activity of agonist and antagonistic muscles to sustain motor adaptations over longer periods (Lavaud et al., 2024).

While previous studies utilized stimulation protocols primarily targeting nociceptive pathways, we aimed to determine whether non-nociceptive sensory feedback, such as low-threshold mechanoreceptive feedback, can also induce sustained motor adaptations. Given that cutaneous sensory feedback plays a major role in mediating locomotor adaptations and recovery after spinal cord injury (Bouyer and Rossignol, 2003; Sławińska et al., 2012), isolating the role of these low-threshold afferents and downstream spinal circuits in sustained motor adaptations could be useful for improving the training of spinal circuits for recovery after spinal cord injury.

In this study, we specifically investigate the role of dI3 neurons in these sustained motor adaptations in response to low-threshold stimulation. Given that dI5s are primarily located in lamina I and II and are involved in various nociceptive pathways (Koch et al., 2018; Xu et al., 2008), they are much less likely to receive low-threshold cutaneous input that terminates in lamina III-IV (Li et al., 2011). Conversely, dI3s are located in lamina IV-V with dendrites extending to lamina III (Bui et al., 2013; Ozyurt et al., 2025), have been shown to integrate both cutaneous low-threshold mechanoreceptive and proprioceptive inputs (Bui et al., 2013, 2016; Ozyurt et al., 2025), and directly excite motoneurons (Bui et al., 2013; Nasiri et al., 2024). Furthermore, dI3s play a role in dynamic corrective movements (Ozyurt et al., 2025) and are crucial for the recovery of locomotor function after spinal cord injury (Bui et al., 2016). Given that the integration of both proprioceptive and high-threshold cutaneous feedback was necessary for sustained motor adaptations (Lavaud et al., 2024), the known circuitry of dI3s suggests these neurons can mediate sustained motor adaptations via low-threshold sensory input.

To selectively recruit cutaneous afferents, we performed closed-loop electrical stimulation of the saphenous nerve. Using a dual Cre- and Flp-recombinase approach driven by Isl1 and Vglut2, respectively, we express the inhibitory DREADD receptor, hM4Di, in dI3 neurons to selectively silence them during motor adaptation paradigms. We show that the inhibition of dI3 neurons impairs sustained motor adaption induced by the stimulation of a cutaneous nerve at a range of stimulation intensities that putatively recruit low-threshold sensory afferents. These findings highlight the potential for different types of sensory information to produce adaptions to motor output on longer timescales through identified spinal circuits.

## MATERIALS & METHODS

### Animals

All animal protocols and experimental procedures at University of Ottawa were followed with respect to the guidelines of the Canadian Council of Animal Care (CCAC) and have been approved by the University of Ottawa Animal Care Committee (BL-3945). As described previously in Goltash et al. (2025), the dI3 neurons were targeted in mice through Cre and FlpO recombinase expression driven by *Isl1* and *Slc17a6* (Vglut2), respectively. *Isl1^Cre+/-^* mice were crossed with *Slc17a6^FlpO+/+^*mice to drive the expression of Cre and FlpO recombinase in dI3 neurons and Vglut2^+^ nociceptive afferents, forming dI3-driver mice. Two genetically defined groups of transgenic mice were used to express hM4Di in dI3 neurons. First, *Isl1^Cre^*^+/-^; *Slc17a6^FlpO^*^+/+^ (dI3-driver) mice were crossed with *RC::FPDi* dual-recombinase responsive fluorescent/DREADD (hM4Di) mice (The Jackson Laboratory, Strain No.: 029040) containing a *frt*-flanked STOP and *loxP*-flanked mCherry::STOP cassete upstream of an HA-tagged hM4Di receptor, resulting in *Isl1^Cre^*^+/-^; *Slc17a6^FlpO^*^+/+^; *RC::FPDi^DREADD^* (dI3^hM4Di^) mice. For the second group, driver mice were given intraspinal injections of pAAV-nEF-Con/Fon DREADD Gi-mCherry (Addgene viral prep # 177672-AAV8), carrying a Cre and Flp dependent and nEF-driven gene for hM4D(Gi) receptor with an mCherry reporter. We refer to these mice as dI3*^AAV-hM4Di^*.

### Intraspinal Injections

Bilateral intraspinal injections were performed to deliver AAV vectors into the lumbar spinal cord. Five-week-old mice were given buprenorphine HCl (0.1 mg/kg, subcutaneously) for preoperative analgesia one hour prior to surgery. Anaesthesia was induced and maintained during surgery using isoflurane (4% induction, 1–2% maintenance in oxygen). Mice were positioned in a stereotaxic frame, and a dorsal midline incision was made over the thoracolumbar vertebrae, and the muscle tissue was retracted to expose the vertebral column. Interlaminar windows were identified between the T11–T12, T12–T13, and T13–L1 vertebrae, corresponding to the approximate location of the lumbar spinal cord. A quartz micropipette (pulled to a ∼30–50 µm tip diameter using a Sutter Instruments P-2000) was attached to a removable-needle Hamilton syringe using a compression fitting and backfilled with paraffin oil. The vector solution was drawn into the syringe by placing the needle tip into a drop of the vector solution and withdrawing at a rate of 1 μL/min using a Harvard Apparatus Pump 11 Elite Nanomite mounted on a micromanipulator. Using the same system, 300 nL of AAV was injected per site at a depth of ∼650-700 µm from the dorsal surface of the spinal cord at a rate of 100 nL/min. Injections were made bilaterally at each interlaminar window, for a total of six injection sites per animal. The micropipette was left in place for 1–2 minutes post-injection to minimize backflow before being slowly withdrawn. Muscle and skin were sutured in layers, and animals were buprenorphine (0.1 mg/kg, Ceva Animal Health) for two days to manage pain and antibiotics (Baytril, Elanco) via drinking water for seven days to prevent infection. Mice were allowed to recover for at least two weeks prior to undergoing spinal cord transection surgery. This surgical protocol was adapted in part from Goltash et al. (2025).

### Chemogenetic Inhibition of dI3 Activity

hM4Di were activated by administration of the selective agonist JHU37160 dihydrochloride (Hello Bio). JHU37160 was dissolved in sterile saline and administered via subcutaneous injection at a dose of 0.5 mg/kg, 10 minutes prior to the start of behavioral testing. To allow full drug washout, a minimum interval of 48 hours was maintained between treatment sessions. EMG and video analysis were done without prior knowledge of the treatment, saline or JHU37160, received.

### Electrode Fabrication

Custom electromyography (EMG) muscle electrodes and nerve cuffs were fabricated using established protocols and commercially available materials and following the protocols described in previous mouse studies (Pearson et al., 2005; Akay et al., 2006; Mayer and Akay, 2018). All electrode assemblies were sterilized by immersion in 3% hydrogen peroxide for 25 minutes, followed by thorough rinsing with sterile distilled water immediately before implantation. Electrodes were tested for continuity and impedance before use to ensure proper function. Paired-wire bipolar EMG electrodes were constructed from 7-strand stainless steel wire (A-M Systems, cat. no. 793200; bare diameter 0.001“, coated diameter 0.0055”). Each electrode was cut to a length of 7 cm from the connector to the needle. The wires were stripped of insulation, twisted together, and knotted at the midpoint to form the recording site. A 27G needle was attached to one end of the electrode pair to facilitate insertion into muscle tissue, while the opposite ends were soldered to gold pins (DigiKey, part no. SAM1155-08-ND) for connection to the headpiece. Nerve cuffs were fabricated using paired stainless-steel wires prepared as above. One end of the wires was threaded through a segment of silicone tubing (Alliedsil; internal diameter 0.020“, outer diameter 0.037”), which served as the cuff. The cuff was designed to be placed around the saphenous nerve and secured with a silk suture passed through the silicone. Both muscle and nerve cuff electrodes were connected to a central connector (DigiKey, part no. SAM15648-ND; length 10 mm), which was inserted subcutaneously in the back of the mouse at the thoracic region. The plastic housing of the connector was coated in a thin layer of biocompatible 5-Minute Epoxy Gel (Devcon) to ensure a smooth, rounded finish and minimize tissue irritation. The epoxy was allowed to cure for at least 24 hours prior to use.

### Spinal Transection

Mice were given analgesia preoperatively via bupivacaine injection at a standard dose. Mice were then anesthetized with 2% isoflurane and placed in a prone position on a heating pad to maintain body temperature during surgery. The skin was incised parallel to the body along the midline of the back, extending from the caudal cervical vertebrae to the caudal thoracic vertebrae. The underlying fascia was removed with surgical forceps, and the muscle attached to the lamina at the thoracic level was dissected away bilaterally from the lamina. The T7 and T8 laminae were cut on both sides with surgical scissors to expose the T10 spinal segment, accounting for the anatomical offset between spinal segments and vertebral levels. Two cuts were made at the T10 level, and the spinal cord segment was removed. Hemostasis was achieved by applying sterile surgical cotton swabs to absorb blood for approximately five minutes. Complete transection was ensured by using a surgical hook to gently scrape horizontally along the spinal column, severing any remaining tissue bridges. The area was again blotted for an additional five minutes until bleeding subsided. Completeness of the transection was confirmed intraoperatively by visual inspection, with particular attention to the lateral aspects of the column to ensure no residual connections remained. Sterile surgical foam (Surgifoam, Johnson & Johnson Medtech) was then placed in the cavity to prevent axonal regeneration across the lesion. The overlying muscle was closed using continuous 6-0 vicryl sutures (Ethicon), and the skin was closed with wound clips, which were removed one week later. Postoperative pain management consisted of subcutaneous buprenorphine (0.1 mg/kg, Ceva Animal Health) administered for three days. To ensure no regeneration occurred during the experimental timeline, post-mortem inspection of the lesion site was performed during the spinal cord dissection.

### Implantations

Following a one-week recovery period, mice received buprenorphine HCl (0.1 mg/kg, subcutaneously) one hour prior to surgery, and anesthesia was induced and maintained with isoflurane throughout the procedure. Two bipolar EMG electrodes were implanted per mouse, targeting the ankle flexor (tibialis anterior, TA) and extensor (gastrocnemius, GS) muscles of the left hindlimb. Additionally, a nerve cuff electrode was placed around the saphenous nerve (SN) of the left hindlimb. The surgical procedures for electrode implantation were followed using established protocols for hindlimb EMG recordings in mice (Akay et al., 2006; Akay, 2014). Incisions were made in the upper neck region, hip, and over the hindlimbs to expose the target muscles and the saphenous nerve. Electrode wires were threaded subcutaneously from the neck incision through the hip incision and down to the hindlimb, then inserted into the muscle tissue parallel to the muscle fibers. The nerve cuff electrode was positioned around the saphenous nerve to enable electrical stimulation of cutaneous afferent fibers supplying the anterior distal hindlimb. A connector was secured subcutaneously in the dorsal thoracic region using 5-0 Prolene sutures, with four interrupted knots anchoring the connector at each corner. All incisions were closed using 7-0 Prolene sutures via an interrupted suture technique with multiple double knots per suture. Each hindlimb incision received 5–6 knots, and the hip incision was similarly closed after feeding the wires through. Postoperative analgesia included buprenorphine (0.1 mg/kg, subcutaneously) administered 6 hours after surgery and then every 12 hours for 48 hours. A topical application of bupivacaine was applied to all incisions at the time of surgery and at each analgesic time point to minimize irritation. For infection prophylaxis, a broad-spectrum antibiotic (Baytril, Elanco) was provided ad libitum in the drinking water for one-week post-surgery. Mice were individually housed in cages placed on a heating pad for the first two days after surgery and then returned to their standard housing conditions. All animals were given at least one week to recover before beginning any acclimation or behavioral testing.

### Recordings and Stimulation

Muscle activity was recorded using a Differential AC Amplifier (A-M Systems, Model 1700) set to a gain of ×1000, with a low cut-off frequency of 300 Hz, a high cut-off frequency of 500 Hz, and the notch filter engaged. Signals were digitized using a Digidata 1440a system (Axon CNS, Molecular Devices) at a sampling rate of 10,000 Hz. Data were acquired using AxoScope software and analyzed with custom Python scripts and GraphPad Prism. Electrical stimulation of the saphenous nerve was performed using a GRASS S88 Stimulator connected to a photoelectric stimulus isolation unit delivering constant current output. The polarity was set to normal, with a current range of 0.1–1.5 mA. Stimulation parameters included a pulse duration of 9 ms and current amplitudes of 0.3, 0.6, or 1.2 mA, set by adjusting the stimulator voltage and confirmed with a multimeter at the isolator output. A 1-second interval was maintained between pulses. At the start of experimentation for each mouse, the stimulation threshold was empirically determined to ensure reliable, selective activation of Aβ sensory fibers, while minimizing recruitment of higher-threshold nociceptive afferents. Threshold was operationally defined as the minimum current amplitude required to elicit a clear, reproducible muscle twitch. This behavioral response was verified in real time by visual inspection of the limb and corroborated by detection of corresponding EMG activity in the target muscle. The current at which a visible limb reflex was reliably evoked in all trials (10/10 responses) was adopted as the animal’s stimulation threshold. Across all mice tested, a current amplitude of approximately 0.3 mA consistently elicited the threshold behavioral response and was thus used for all subsequent closed-loop trials. Threshold determinations were not blinded to animal or treatment. They were determined following saline injections to ensure standardized and reproducible stimulation conditions across the study.

### Tissue Collection & Immunohistochemistry

Following dissection, each cord was fixed by immersion in 5 mL of 4% paraformaldehyde (PFA) for 16 hours at 4°C. After fixation, tissues were transferred to a 30% sucrose solution in PBS for cryoprotection, then embedded in OCT compound and stored at –80°C until sectioning. Cervical and lumbar segments of the spinal cord were sectioned at a thickness of 25 μm using a cryostat, and sections were stored at –4°C until staining. For immunohistochemistry, sections were mounted onto standard glass slides and rinsed in PBS. Blocking was performed using 5% goat serum in PBS for 30 minutes at room temperature. To enhance detection of hM4Di-mCherry, sections were incubated with a rabbit polyclonal anti-RFP antibody (Rockland, 600-401-379-RTU; 1:500 dilution) for 48 hours at 4°C. After washing, sections were incubated with a goat anti-rabbit HRP-conjugated secondary antibody (Abcam, ab7090; 1:2000 dilution) for 3 hours at room temperature. Signal amplification was achieved using tyramide signal amplification (TSA) with 690 Opal dye (1:150 dilution in TSA buffer) for 8 minutes, followed by several washes in PBS. Nuclei were counterstained with DAPI (1:500 dilution) for 3 hours, included during secondary antibody incubation. After staining, sections were coverslipped with Immunomount. Confocal imaging was performed using an Olympus FV1000 BX61 LSM microscope. Images were processed and analyzed using ImageJ software.

### Closed-loop Stimulation

To investigate spinal motor learning, we implemented a closed-loop feedback system in which electrical stimulation was delivered contingently on the mouse’s real-time toe position. Each mouse was placed in a custom-made cardboard harness with a Velcro strap, individually fitted to ensure security without restricting breathing. The harness was suspended above the ground using a repurposed horizontal steel arm, allowing the left hindlimb to move freely. The mouse’s torso was held horizontally, with the tail taped above the camera’s field of view. To reduce stress and movement, the mouse’s front paws rested on a large tube positioned in front of the animal. High-speed video was captured using a BASLER acA640-750µm camera (100 frames per second, 640×480 pixels), positioned directly in front of the left hindlimb. The right hindpaw was masked with black velvet tape to prevent mis-tracking, and black velvet paper was used as a backdrop to enhance contrast. Three LED lights illuminated the hindlimb for optimal video clarity. Video acquisition and real-time pose estimation were performed using a custom Python script built on DeepLabCut-Live version 1.0.2. The neural network was trained on 800 manually labeled frames from practice session videos. DeepLabCut-Live pose estimation inferences of toe and threshold vertical positions were performed on an Intel Core i9-13900K CPU and RTX 4070TI GPU, achieving 10-20 ms per frame. After harnessing, each mouse was acclimated for 10 minutes to allow the hindlimb to settle into its natural resting position. A ruler visible in the camera’s field of view enabled spatial calibration. The script tracked both the toe and a visible vertical threshold using DeepLabCut-Live. The vertical threshold was set approximately 2 mm above the resting toe position and could be adjusted manually by turning a knob. When the tracked toe dropped below the threshold, the Python script instructed a microcontroller (Arduino UNO, ELEGOO UNO R3) to send a TTL pulse. This pulse was routed through a custom “TTL to contact closure” converter, which triggered the GRASS S88 stimulator to deliver a stimulation pulse to the SN. The measured latency from threshold crossing to stimulation delivery was 10-20 ms. To ensure accurate triggering and minimize false positives, the script displayed the video feed and tracking confidence in real time, only allowing stimulation when DeepLabCut-Live’s confidence exceeded 90%. A 1-second grace period was enforced between stimulations to limit frequency. For the 1.2 mA experiment, a learner and a control mouse were used. Both the learner and the control mice received closed-loop stimulation conditional on the toe of the learner mouse crossing the threshold. The toe height of the control mouse had no bearing on stimulation. Both mice were harnessed and positioned identically. All experimental data such as video, estimated toe-to-threshold distances, tracked positions, and tracking confidence were recorded by the custom Python script.

### Data & Statistical Analysis

Gastrocnemius (GS) and tibialis anterior (TA) muscle activity was analyzed by applying a root-mean-square (RMS) envelope of the EMG signal using a continuous moving window with a duration of 20 ms. This step produced a smooth representation of muscle activation amplitude across time. Manual annotation was then performed using a custom-built Python script. All annotations were performed blind to both animal identity and treatment condition (e.g., saline vs. JHU37160). Stimulation artifacts were qualitatively distinguishable from genuine muscle activity and excluded from analysis. For each animal, results were averaged across all available trials under each treatment condition (saline or JHU37160), yielding a per-animal mean for each metric and condition. GraphPad Prism 8.0.2 and custom Python script was used for statistical analysis and production of graphs. Data are reported as mean ± S.D and the significance level was set at p < 0.05. Comparisons were made using either unpaired t-test (two-tailed), paired t-tests (two-tailed), or repeated measures one-way ANOVA followed by comparison between pairs of groups using Tukey’s or Sidak’s multiple comparison test. \**p* < 0.05, \*\**p* < 0.01, \*\*\**p* < 0.001.

## RESULTS

### Closed-loop stimulation of saphenous nerve induces spinal motor adaptation

We first attempted to generate motor adaptation in spinalized mice using a closed-loop stimulation approach whereby toe height below a threshold level triggers stimulation of the left saphenous nerve. Prior to performing this closed-loop paradigm, we allowed the mice to recover for 7 days after the spinal transection and another 7 days after implantation of the nerve cuff electrodes (**Fig. 1*A***). To confirm a previous study that stimulation contingent on limb position is required for eliciting this motor adaptation (Lavaud et al., 2024), a “learner” and a “control” mouse were paired such that the left saphenous nerves of both animals were stimulated simultaneously whenever the toe of the learner mouse fell below a target (2 mm from resting position) (**Fig. 1*B***). Previous studies investigating motor adaptation at the spinal level by electrical stimulation implicated high-threshold mechanoreceptive or nociceptive inputs (Buerger and Fennessy, 1971; Grau, 2014; Lavaud et al., 2024). To recruit these fibers, we stimulated the saphenous nerve at 4 times the threshold for visible motor reflexes (Koga et al., 2005).

**Figure 1.**
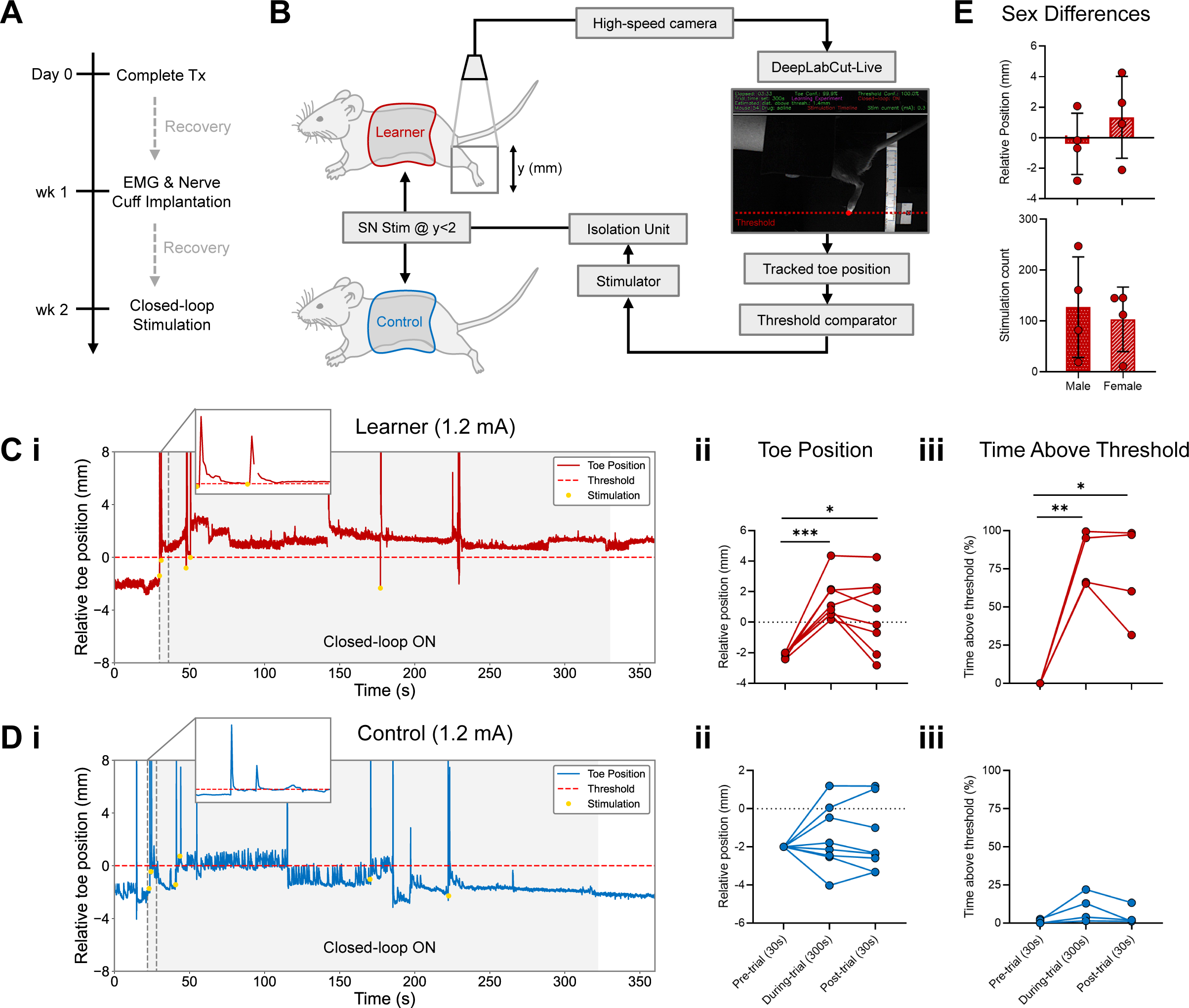
Closed-loop stimulation of saphenous nerve induces spinal motor adaptation. ***A.*** Schematic of the experimental timeline. Mice underwent complete spinal transection at approximately thoracic level T10 on Day 0. On Day 7, lateral gastrocnemius (GS) and tibialis anterior (TA) EMG electrodes were implanted, and a saphenous nerve (SN) cuff was implanted. Mice were allowed to recover for at least 7 days before experimentation. ***B.*** Diagram of the closed-loop electrical stimulation procedure applied to learner and control mice. Learner mice receive stimulation contingent on their limb position. Control mice receive the same stimulations as Learner mice; thus, stimulations applied to control mice are not contingent on their limb position. ***C.*** (***i***) Representative trace of toe position during a closed-loop stimulation trial (saline administration with 1.2 mA stimulations). Stimulation times and relative toe position at the time of stimulation is depicted by yellow dots. (***ii***) Average toe vertical position across trial phases in closed-loop stimulation trials (saline, 1.2 mA; *n* = 8). Pre-trial vs during-trial, *p* = 0.0004; pre-trial vs post-trial, *p* = 0.0427; during-trial vs post-trial, *p* = 0.2061. (***iii***) Proportion of time the toe remained above threshold during each trial phase with saline and 1.2 mA stimulations (*n* = 4). Pre-trial vs during-trial, *p* = 0.0061; pre-trial vs post-trial, *p* = 0.0419; during-trial vs post-trial, *p* = 0.5382. ***D.*** (***i***) Control stimulation trial paired with the trial shown in panel C. The same stimulation times were delivered to this subject, but independently of toe position, administered with saline. (***ii***) Average vertical toe position across trial phases in control stimulation trials (saline, 1.2 mA; *n* = 8). Pre-trial vs during-trial, *p* = 0.7057; pre-trial vs post-trial, *p* = 0.8006; during-trial vs post-trial, *p* = 0.9493. (***iii***) Proportion of time the toe remained above threshold during each trial phase (saline, 1.2 mA; *n* = 4). Pre-trial vs during-trial, *p* = 0.0830; pre-trial vs post-trial, *p* = 0.5411; during-trial vs post-trial, *p* = 0.1205. ***E.*** Post-trial toe position (top, *p* = 0.3388) and stimulation count of each trial (bottom, *p* = 0.6980) and for male (*n* = 4) and female (*n* = 4) mice. Each point represents the mean value for a single animal, and each line connects the mean values from the same subject. Statistical analysis, repeated-measures one-way ANOVA with Tukey’s multiple comparisons test and two-tailed unpaired t test. \**p* < 0.05; \*\**p* < 0.01; \*\*\**p* < 0.001.

The learner mice exhibited sustained periods of elevated toe positioning (**Fig. 1*Ci***). In contrast, control mice failed to maintain an elevated toe position during the trial (**Fig. 1*Di***). We analyzed the average toe position and the relative time spent above the threshold for three epochs: 30s before turning on the closed-loop stimulation, during five minutes of closed-loop stimulation, and 30s after removing terminating the closed-loop stimulation. In learner mice, the average toe position was significantly higher during the five minutes of closed-loop stimulation (Pre-trial vs During-trial: *n* = 8 mice; *p* = 0.0004) and during the 30s post-closed loop stimulation epoch (Pre-trial vs Post-trial: *n* = 8 mice; *p* = 0.0427) relative to the 30s epoch before turning on the closed-loop stimulation (**Fig. 1*Cii***). Conversely, there were no differences in toe elevation in the three epochs for control mice (*n* = 8; all *p* > 0.7057) (**Fig. 1*Dii***). Similarly, in learner mice the time spent above threshold, measured in percentage of the total duration of each epoch, was greater during the 5min of closed-loop stimulation (Pre-trial vs During-trial; *n* = 4; *p* = 0.0061) and the 30s post-closed loop stimulation epoch (Pre-trial vs Post-trial; *n* = 4; *p* = 0.0419) relative to the toe position pre-trial; however, there were no differences between the three epochs in control mice (*n* = 4; all *p* > 0.0830) (**Fig. 1*Ciii, Diii***). These results demonstrate that closed-loop stimulation of the left saphenous nerve contingent on limb position induces sustained adaptation of limb positioning, as evidenced by sustained elevation of toe position.

### Sex does not influence closed-loop motor learning metrics

Previous research has demonstrated that males and females can exhibit significant differences in pain perception and processing, with females often displaying greater pain sensitivity (Fillingim et al., 2009; Mogil, 2012). Given our 4x threshold stimulation at 1.2 mA is expected to robustly recruit both Aδ and C fibers (Kendroud et al., 2025), we assessed whether sex influenced behavioural outcomes in this closed-loop learning paradigm. There were no significant differences between males and females in any of the primary outcome measures. We observed no differences in post-trial toe position or stimulation count per trial (*n* = 4 males, *n* = 4 females; *p* = 0.3388 and *p* = 0.6980 respectively) (**Fig. 1*E***). These findings indicate that, despite well-established sex differences in pain processing, sex did not significantly affect the number of stimulations required and the persistence of learned hindlimb elevation.

### Silencing of dI3 neurons limits motor adaptation at 1.2 mA stimulation

We next asked whether dI3 neurons were involved in the form of motor adaptation generated above. Using an *Isl1^Cre^*^+/-^; *Slc17a6^FlpO^*^+/+^ mouse line to drive Cre and FlpO expression in dI3 neurons, we expressed the inhibitory DREADD receptor hM4Di in these neurons through transgenic recombination (**Fig. 2*Ai***; *n* = 4) or through an AAV-mediated conditional expression (**Fig. 2*Aii, B***; *n* = 4). For the AAV driven expression of hM4Di, we performed intraspinal injections of pAAV-nEF-Con/Fon DREADD Gi-mCherry 2 weeks prior to the spinal transection surgeries, followed by a similar timeline for nerve cuff electrode implants and stimulations (**Fig. 2*C***). Immunofluorescence images of lumbar spinal cord in dI3^AAV-hM4Di^ mice revealed hM4Di-mCherry-positive cells in both the superficial dorsal horn (laminae I-II) and the deeper dorsal/intermediate laminae (laminae V-VII) (**Fig. 2*D***). Although we did not directly label dI3s, previous work has demonstrated that they are located predominantly in laminae V-VII of the lumbar spinal cord (Bui et al., 2013). Based on this, the hM4Di-mCherry-positive cells present in these laminae are likely to correspond to dI3s (**Fig. 2*D***). In addition, a subset of sensory afferents involved in mechanical nociception also express Isl1 and Vglut2 (Lagerström et al., 2010; Goltash et al., 2025). Afferents mediating mechanical nociception, including Aδ and C fibers, are known to terminate primarily in the superficial dorsal horn, specifically within laminae I–II (Light and Perl, 1979; Pan and Pan, 2004; Todd, 2017). Therefore, the hM4Di-mCherry–positive signal observed in laminae I–II likely represents the terminations of these nociceptive afferents (**Fig. *2D***).

**Figure 2.**
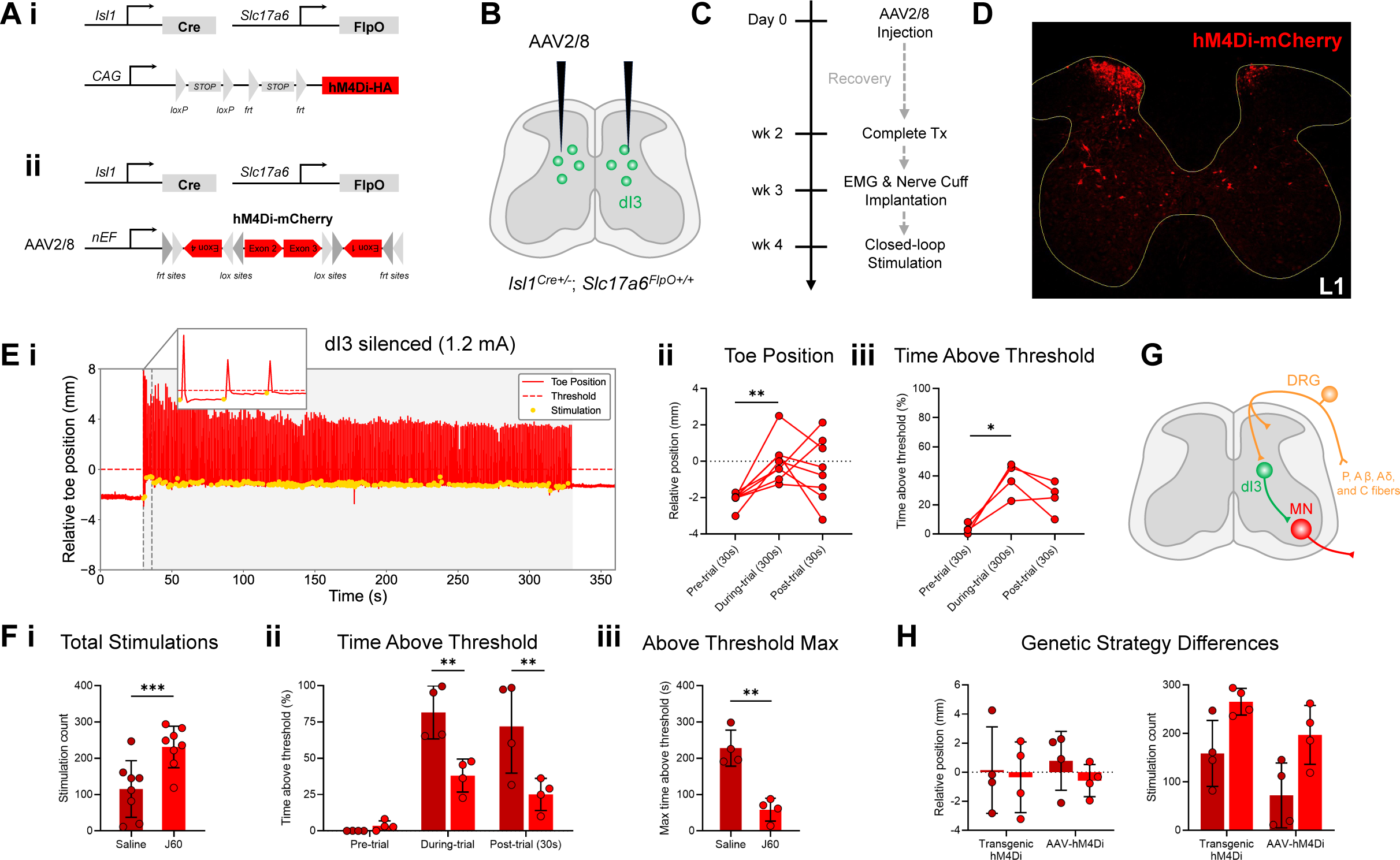
Silencing of dI3 neurons limits motor adaptation at 1.2 mA stimulation. ***A.*** The dI3 neurons were targeted in mice through Cre and FlpO recombinase expression driven by *Isl1* and *Slc17a6* (Vglut2), respectively. Two genetically defined groups of transgenic mice were used to express hM4Di in dI3 neurons. (***i***) First, dI3-driver mice were crossed with *RC::FPDi* dual-recombinase responsive fluorescent/DREADD (hM4Di) mice. (***ii***) For the second group, driver mice were given intraspinal injections of pAAV-nEF-Con/Fon DREADD Gi-mCherry, carrying a Cre and Flp dependent and nEF-driven gene for hM4D(Gi) receptor with an mCherry reporter (dI3^AAV-hM4Di^). ***B.*** Injections were made bilaterally at each interlaminar window for a total of six injection sites per animal, roughly targeting L1-L5 segments of the spinal cord. ***C.*** Schematic of the experimental timeline. dI3^AAV-^hM4Di mice received intraspinal injections of AAV2/8 at Day 0 and were allowed to recover for at least 2 weeks before complete transection surgery. Mice underwent complete spinal transection at approximately thoracic level T10 at week 2. On week 3, lateral gastrocnemius (GS) and tibialis anterior (TA) EMG electrodes were implanted, and a saphenous nerve (SN) cuff was implanted. Mice were then allowed to recover for at least 7 days before experimentation. ***D.*** Sample immunofluorescence image of hM4Di-mCherry expression (red) at an L1 spinal cord section from a representative dI3^AAV-hM4Di^ mouse. ***E.*** (***i***) Representative trace of toe position during a closed-loop stimulation trial (JHU37160 administration with 1.2 mA stimulations). Yellow dots mark the toe position at the time of stimulation. (***ii***) Average toe vertical position across trial phases in closed-loop stimulation trials (JHU37160, 1.2 mA; *n* = 8). Pre-trial vs during-trial, *p* = 0.0047; pre-trial vs post-trial, *p* = 0.0864; during-trial vs post-trial, *p* = 0.7880. (***iii***) Proportion of time the toe remained above threshold during each trial phase with JHU37160 and 1.2 mA stimulations (*n* = 4). Pre-trial vs during-trial, *p* = 0.0252; pre-trial vs post-trial, *p* = 0.1036; during-trial vs post-trial, *p* = 0.2353. ***F.*** (***i***) Stimulation count per trial at 1.2 mA stimulation (*n* = 8). Each point represents the mean number of stimulations an animal received in one trial (300s) averaged across multiple trials. Saline vs JHU37160, *p* = 0.0001. (***ii***) Proportion of time the toe remained above threshold during each trial phase with 1.2 mA stimulation (*n* = 8). Each point represents the mean value for a single animal, averaged across multiple trials. Saline vs JHU37160 (pre-trial), *p* = 0.9653; saline vs JHU37160 (during-trial), *p* = 0.0048; saline vs JHU37160 (post-trial), *p* = 0.0033. (***iii***) Longest continuous duration that the toe was held above threshold with 1.2 mA stimulation (*n* = 8). Each point represents the maximum value that an animal held the toe above the threshold across any trial. Saline vs JHU37160, *p* = 0.0044. ***G.*** Schematic of the role of low and high-threshold sensory afferents and dI3 neurons in motor adaptation. DRG: dorsal root ganglion, MN: motoneuron, P: proprioceptive. ***H.*** Differences in post-trial toe position (left, transgenic hM4Di vs AAV-hM4Di: saline treated: *p* = 0.9050; J60 treated: *p* = 0.9870) and stimulation count (right, transgenic hM4Di vs AAV-hM4Di: saline treated: *p* = 0.1113; J60 treated: *p* = 0.2299) between transgenic hM4Di (*n* = 4) and AAV-hM4Di (*n* = 4) mice. Each point represents the mean stimulation count for each animal, averaged across multiple trials. The saline and JHU37160 groups include the same animals. Statistical analysis, Repeated-measures two-way ANOVA with Sidak’s multiple comparisons test or two-tailed, paired t-test. \**p* < 0.05, \*\**p* < 0.01, \*\*\*\**p* < 0.001, \*\*\*\**p* < 0.0001.

We conducted closed-loop and control trials at 4x threshold (1.2 mA) following JHU37160 administration. In closed-loop trials with dI3-silenced (**Fig. 2*Ei***), animals exhibited a significant increase, but subthreshold average toe position during the stimulation period compared to the pre-trial baseline (Pre-trial vs During-trial: *n* = 8; *p* = 0.0047), suggesting that some response occurred (**Fig. 2*Eii***). However, this improvement did not persist after stimulation ceased, as post-trial toe position was not significantly different from baseline (Pre-trial vs Post-trial: *n* = 8; *p* = 0.0864) (**Fig. 2*Eii***). A similar pattern was observed for the proportion of time the toe remained above threshold (**Fig. 2E*iii***; *n* = 4; Pre-trial vs During-trial: *p* = 0.0252; Pre-trial vs Post-trial: *p* = 0.1036). These findings suggest that although toe position increased during the stimulation period, the effects were short-lived and did not consolidate into longer-lasting post-trial changes. This pattern may reflect a transient or less stable form of motor adaptation that emerges during stimulation but fails to persist once stimulation ends. Another possibility is the acute, reflexive response to strong stimulation rather than true sustained adaptation that causes the toe to be above the threshold during the trial (**Fig. 2*Ei***).

To further evaluate differences in generating motor adaptation in our closed-loop paradigm with and without dI3 neuron activity, we assessed three key parameters between saline and JHU37160 administered groups: the total number of stimulations delivered during each 5min trial (stimulation count), the proportion of time the toe was maintained above the threshold (above-threshold time), and the longest continuous duration the toe was held above threshold during any trial for each animal. A lower stimulation count suggests more effective learning since stimulations are only delivered when the toe is below the threshold. Given that our stimulations were delivered at 1 Hz, the maximum possible number of stimulations per trial was 300. Saline-treated mice required significantly fewer stimulations than JHU37160-treated mice (**Fig. *2Fi***, *n* = 8; *p* = 0.0001). Similarly, the proportion of time the toe was maintained above threshold during the trial was significantly greater with saline than with JHU37160 (**Fig. 2*Fii***, *n* = 8; *p* = 0.0048), and this effect persisted into the post-trial period (**Fig. 2*Fii***; *n* = 8; *p* = 0.0033). For the longest continuous duration the toe was held above threshold, saline-treated mice outperformed JHU37160-treated mice (**Fig. 2*Fiii***, *n* = 8; *p* = 0.0044).

Overall, our findings suggest that closed-loop stimulation of nociceptive and cutaneous afferents and spinal dI3 neurons are involved in mediating these spinal motor adaptations (**Fig. 2*G***). It is important to note that hM4Di expression in some Aδ and C fibers (Goltash et al., 2025) means these nociceptive afferents will also be hyperpolarized upon hM4Di activation (**Fig. 2*G***). With the aim to isolate the function of dI3s, we carefully control the stimulation parameters later in this study to avoid confounding activation of nociceptive fibers.

### Genetic strategy of hM4Di expression does not influence motor learning metrics

Given that a transgenic strategy is likely to induce more broad expression of hM4Di compared to bilateral AAV injections at three lumbar segments, we investigated whether the method of expression of hM4Di receptors in Isl1^+^/Vglut2^+^ cells (dI3^hM4Di^ and dI3^AAV-hM4Di^) influences motor adaptation. We compared the two groups during closed-loop trials at 1.2 mA with saline or JHU37160 treatment (**Fig. 2*H***; *n* = 4 dI3^hM4Di^; *n* = 4 dI3^AAV-hM4Di^). Relative post-trial toe position did not differ between dI3^hM4Di^ and dI3^AAV-hM4Di^ with either saline (*n* = 4 per group; *p* = 0.9050) or JHU37160 treatment (*n* = 4 per group; *p* = 9870) (**Fig. 2*H***). Total stimulation count also did not differ between dI3^hM4Di^ and dI3^AAV-hM4Di^ mice with either saline (*n* = 4 per group; *p* = 0.1113) or JHU37160 treatment (*n* **=** 4 per group; *p* = 0.2299) (**Fig. 2*H***). Overall, these findings suggest that the method used to express hM4Di receptors in Isl1^+^/Vglut2^+^ cells does not significantly impact motor adaptation metrics during closed-loop trials with 1.2 mA stimulations.

### Silencing of dI3 neurons limits motor adaptation at 0.6 mA stimulation

Given that our objective is to determine the role of dI3s in motor adaptation using a closed-loop stimulation paradigm through LTMR input, we next performed these closed-loop stimulations at lower thresholds to avoid activation of nociceptive fibres in the saphenous nerve. Since the control stimulation experiment with 1.2 mA already confirmed that learning does depend on the contingency of the stimulation, we did not utilize control mice for lower intensity stimulations. A previous study in rats has shown that the activation threshold for C fibers is approximately 4 times higher, and for Aδ fibers approximately 2.5 times higher, than that of Aβ fibers (Koga et al., 2005). Using our saphenous nerve cuff, we determined the behavioral threshold to be approximately 0.3 mA amongst animals tested. Accordingly, we conducted closed-loop stimulation trials at 0.6 mA (2x threshold) in saline and JHU37160 administered mice to selectively activate a large portion of Aβ fibers.

Saline animals demonstrated elevated toe positioning above threshold during the closed-loop stimulation period, with this effect persisting into the post-trial phase (**Fig. 3*Ai***). In contrast, mice with silenced dI3s failed to maintain toe elevation during or post-trial (**Fig. 3*Bi***). As previously done, we quantified the average toe position and proportion of time with toe above threshold at three epochs: pre-trial, during-trial, and post-trial. In saline mice, we found that the average toe position was also significantly elevated during (Pre-trial vs During-trial: *n* = 4; *p* = 0.0081) and after the trial (Pre-trial vs Post-trial: *n* = 4; *p* = 0.0229) relative to pre-trial; however, when dI3s were silenced, the average toe positioned remained unaltered during or post-trial (*n* = 4; *p* > 0.05) (**Fig. 3*Aii**, Bii***). Similarly, the proportion of time above threshold was increased during (Pre-trial vs During-trial: *n* = 4; *p* < 0.0001) and after the trial (Pre-trial vs Post-trial: *n* = 4; *p* = 0.0014) compared to before the onset of closed-loop stimulation; however, dI3 inhibition led to no significant changes to the proportion of time the toe remained above threshold (*n* = 4; *p* > 0.05) (**Fig. 3*Aiii**, Biii***).

**Figure 3.**
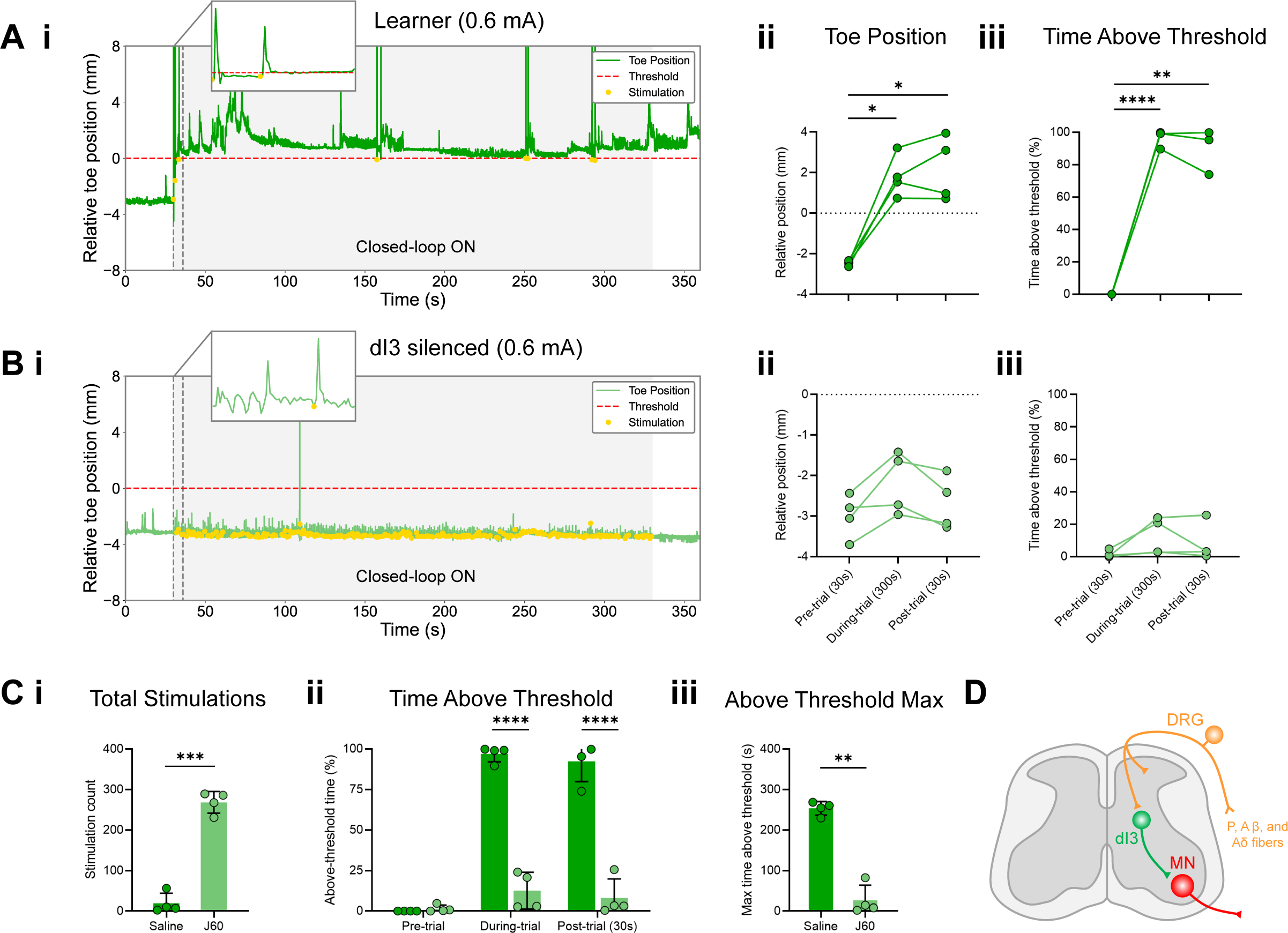
Silencing of dI3 neurons limits motor adaptation at 0.6 mA stimulation. ***A.*** (***i***) Representative trace of toe position during a closed-loop stimulation trial at 0.6 mA stimulations with saline administration. Yellow dots mark toe position at the time of stimulation. (***ii***) Average vertical toe position across trial phases in closed-loop stimulation trials (saline, 0.6 mA; *n* = 4). Pre-trial vs during-trial, *p* = 0.0081; pre-trial vs post-trial, *p* = 0.0229; during-trial vs post-trial, *p* = 0.6838. (***iii***) Proportion of time the toe remained above threshold during each trial phase with 0.6 mA stimulations (*n* = 4). Pre-trial vs during-trial, *p* < 0.0001; pre-trial vs post-trial, *p* = 0.0014; during-trial vs post-trial, *p* = 0.5423. ***B.*** (***i***) Representative trace of toe position during a closed-loop stimulation trial at 0.6 mA stimulations with JHU37160 administration. (***ii***) Average toe vertical position across trial phases in closed-loop stimulation trials (JHU37160, 0.6 mA; *n* = 4). Pre-trial vs during-trial, *p* = 0.6555; pre-trial vs post-trial, *p* = 0.9991; during-trial vs post-trial, *p* = 0.6856. (***iii***) Proportion of time the toe remained above threshold during each trial phase (JHU37160, 0.6 mA; *n* = 4). Pre-trial vs during-trial, *p* = 0.2365; pre-trial vs post-trial, *p* = 0.5967; during-trial vs post-trial, *p* = 0.6176. ***C.*** (***i***) Stimulation count per trial at 0.6 mA stimulation (*n* = 4). Each point represents the mean number of stimulations an animal received in one trial (300s) averaged across multiple trials. Saline vs JHU37160, *p* = 0.0004. (***ii***) Proportion of time the toe remained above threshold during each trial phase with 0.6 mA stimulation (*n* = 4). Each point represents the mean value for a single animal, averaged across multiple trials. Saline vs JHU37160 (pre-trial), *p* = 0.9798; saline vs JHU37160 (during-trial), *p* < 0.0001; saline vs JHU37160 (post-trial), *p* < 0.0001. (***iii***) Longest continuous duration that the toe was held above threshold with 0.6 mA stimulation (*n* = 4). Each point represents the maximum value that an animal held the toe above the threshold across any trial. Saline vs JHU37160, *p* = 0.0013. The saline and JHU37160 groups include the same animals. Statistical analysis, Repeated-measures two-way ANOVA with Sidak’s multiple comparisons test or two-tailed, paired t-test. \**p* < 0.05, \*\**p* < 0.01, \*\*\**p* < 0.001, \*\*\*\**p* < 0.0001. ***D.*** Schematic of the putative low and high-threshold sensory afferents to dI3 neurons involved in this form of motor adaptation at 0.6 mA stimulation strength. DRG: dorsal root ganglion, MN: motoneuron, P: proprioceptive.

Next, we compared the same three parameters previously mentioned between saline and dI3-silenced groups: the total number of stimulations delivered during each 300s trial (stimulation count), the proportion of time the toe was maintained above the threshold (above-threshold time), and the longest continuous duration the toe was held above threshold during any trial for each animal. We found that saline-treated mice required significantly fewer stimulations than dI3 silenced mice during 0.6 mA stimulation (**Fig. *3Ci***, *n* = 4; *p* = 0.0004). Similarly, the proportion of time the toe was maintained above threshold during and post-trial significantly decreased after dI3 silencing (**Fig. 3*Cii***, *n* = 4; *p* < 0.0001). Furthermore, saline-treated mice outperformed JHU37160-treated mice in the longest continuous duration toe elevation above threshold (**Fig. 3*Ciii***, *n* = 4; *p* = 0.0013). Overall, these results demonstrate that closed-loop stimulation reliably induces spinal motor adaptations with 0.6 mA stimulation intensities, which primarily recruits Aβ fibres and potentially a small portion of Aδ fibres, and that dI3 neurons, which are known to integrate inputs from these afferents, are necessary for mediating these spinal motor adaptations (**Fig. 3*D***).

### Silencing of dI3 neurons limits motor adaptation at 0.3 mA stimulation

We next performed the same experiments at an even lower stimulation intensity, specifically at threshold, to ensure we only recruit Aβ fibres. Again, since the control stimulation experiment with 1.2 mA already confirmed that learning does depend on the contingency of the stimulation, we also did not utilize control mice for these low-threshold stimulations.

Even with 0.3mA stimulations, we observed that saline animals demonstrated elevated toe positioning above threshold during the closed-loop stimulation period and persisted with these toe elevations into the post-trial phase (**Fig. 4*Ai***). In contrast, mice with silenced dI3s failed to maintain toe elevation during or post-trial (**Fig. 4*Bi***). We again quantified the average toe position and proportion of time with toe above threshold at the three epochs: pre-trial, during-trial, and post-trial. In saline mice, we found that the average toe position was also significantly elevated during (Pre-trial vs During-trial: *n* = 4; *p* = 0.0262) and after the trial (Pre-trial vs Post-trial: *n* = 4; *p* = 0.0139) relative to pre-trial; however, dI3 silencing prevented any changes in the average toe position during or post-trial compared to before the trial (*n* = 4; *p* > 0.05) (**Fig. 4*Aii**, Bii***). Similarly, the proportion of time above threshold was increased with 0.3 mA during closed-loop stimulation (Pre-trial vs During-trial: *n* = 4; *p* < 0.0337) and after the trial (Pre-trial vs Post-trial: *n* = 4; *p* = 0.0379) relative to before the onset of closed-loop stimulation; however, dI3 inhibition prevented any significant changes to the proportion of time the toe remained above threshold during and after the trial compared to before the trial (*n* = 4; *p* > 0.05) (**Fig. 4*Aiii**, Biii***).

**Figure 4.**
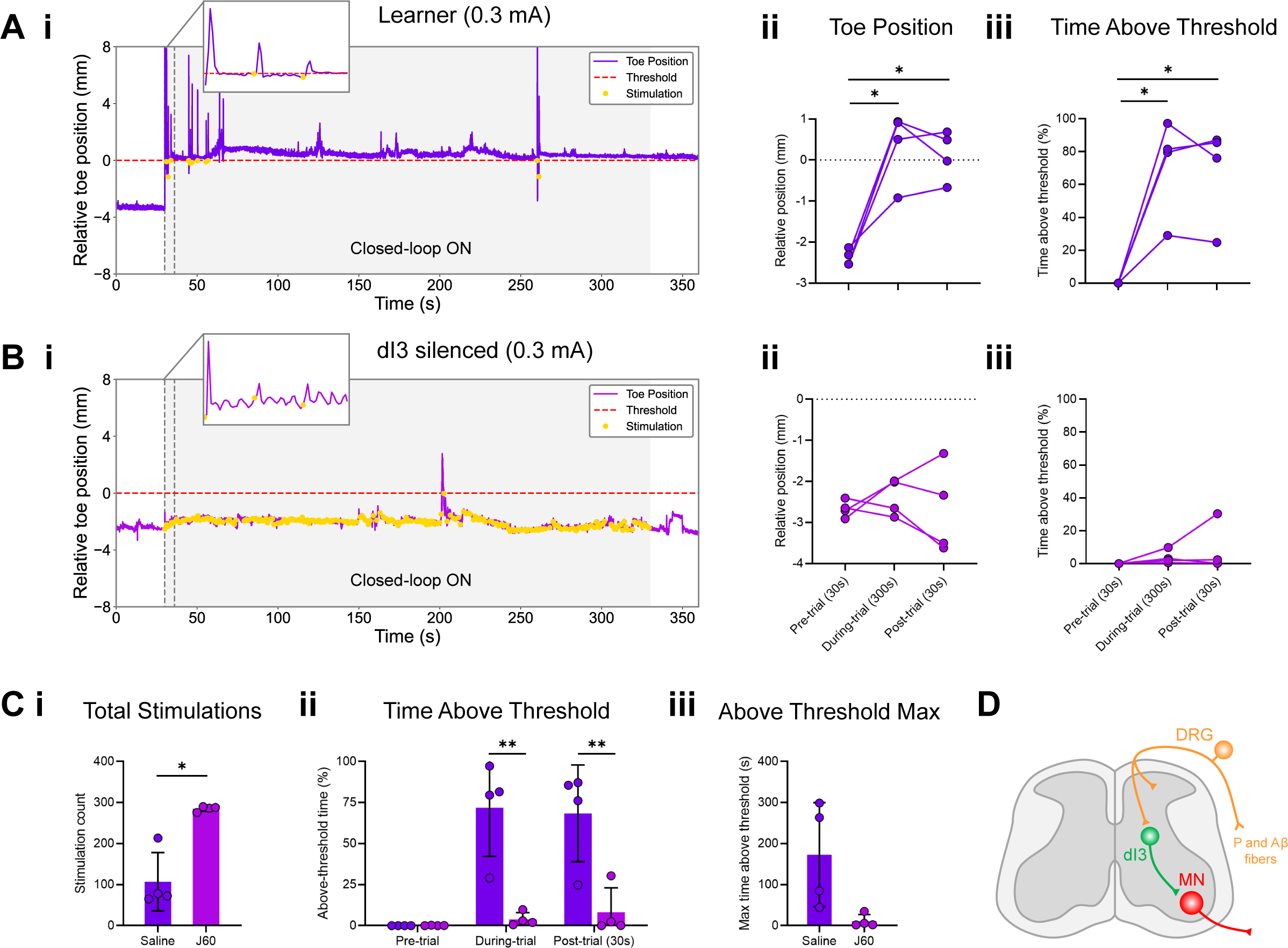
Silencing of dI3 neurons limits motor adaptation at 0.3 mA stimulation. ***A.*** (***i***) Representative trace of toe position during a closed-loop stimulation trial at 0.3 mA stimulations with saline administration. Yellow dots mark toe position at the time of stimulation. (***ii***) Average toe vertical position across trial phases in closed-loop stimulation trials (saline, 0.3 mA; *n* = 4). Pre-trial vs during-trial, *p* = 0.0262; pre-trial vs post-trial, *p* = 0.0139; during-trial vs post-trial, *p* = 0.7037. (***iii***) Proportion of time the toe remained above threshold during each trial phase with 0.3 mA stimulations (*n* = 4). Pre-trial vs during-trial, *p* = 0.0337; pre-trial vs post-trial, *p* = 0.0379; during-trial vs post-trial, *p* = 0.8595. ***B.*** (***i***) Representative trace of toe position during a closed-loop stimulation trial at 0.3 mA stimulations with JHU37160 administration. (***ii***) Average toe vertical position across trial phases in closed-loop stimulation trials (JHU37160, 0.3 mA; *n* = 4). Pre-trial vs during-trial, *p* = 0.6555; pre-trial vs post-trial, *p* = 0.9991; during-trial vs post-trial, *p* = 0.6856. (***iii***) Proportion of time the toe remained above threshold during each trial phase (JHU37160, 0.3 mA; *n* = 4). Pre-trial vs during-trial, *p* = 0.3162; pre-trial vs post-trial, *p* = 0.5805; during-trial vs post-trial, *p* = 0.7238. ***C.*** (***i***) Stimulation count per trial at 0.3 mA stimulation (*n* = 4). Each point represents the mean number of stimulations an animal received in one trial (300s) averaged across multiple trials. Saline vs JHU37160, *p* = 0.0149. (***ii***) Proportion of time the toe remained above threshold during each trial phase with 0.3 mA stimulation (*n* = 4). Each point represents the mean value for a single animal, averaged across multiple trials. Saline vs JHU37160 (pre-trial), *p* > 0.9999; saline vs JHU37160 (during-trial), *p* = 0.0012; saline vs JHU37160 (post-trial), *p* = 0.0024. (***iii***) Longest continuous duration that the toe was held above threshold with 0.3 mA stimulation (*n* = 4). Saline vs JHU37160, *p* = 0.0982. Each point represents the maximum value that an animal held the toe above the threshold across any trial. The saline and JHU37160 groups include the same animals. Statistical analysis, Repeated-measures two-way ANOVA with Sidak’s multiple comparisons test or two-tailed, paired t-test. \**p* < 0.05, \*\**p* < 0.01, \*\*\**p* < 0.001, \*\*\*\**p* < 0.0001. ***D.*** Schematic of the putative low-threshold sensory afferents to dI3 neurons involved in this form of motor adaptation at 0.3 mA stimulation strength. DRG: dorsal root ganglion, MN: motoneuron, P: proprioceptive.

We again compared the same three parameters previously mentioned between saline and JHU37160 administered groups for 0.3 mA stimulations: the total number of stimulations delivered during each 300s trial (stimulation count), the proportion of time the toe was maintained above the threshold (above-threshold time), and the longest continuous duration the toe was held above threshold during any trial for each animal. We found that saline-treated mice required significantly fewer stimulations than dI3 silenced mice (**Fig. *4Ci***, *n* = 4; *p* = 0.0149). Similarly, the proportion of time the toe was maintained above threshold during and post-trial significantly decreased after dI3 silencing (**Fig. 3*Cii***, *n* = 4; During-trial vs Post-trial: *p* < 0.0012; Pre-trial vs Post-trial: *p* = 0.0024). Lastly, we observed no significant difference in the longest continuous duration of toe elevation above threshold for both groups; however, there was a trend towards a decreased performance after dI3 silencing (**Fig. 4*Ciii***, *n* = 4; *p* = 0.0982). These findings strengthen our observations that closed-loop stimulation of LTMRs reliably induces spinal motor adaptations, and dI3 neurons that are known to integrate these sources of sensory input are necessary for mediating these spinal motor adaptations (**Fig. 4*D***).

## DISCUSSION

Sustained adaptation of muscle activity in spinalized settings has been shown in response to electrical stimulation of nociceptive afferents as well as stimulation of muscles and the various afferents that innervate them (Grau, 2014; Lavaud et al., 2024). In these experiments, hindlimbs adapt to context-dependent stimulation by maintaining flexion or elevated toe height for a length of time, outlasting the activation of simple spinal reflexes. Several classes of spinal neurons have been linked to this form of motor adaptation (Lavaud et al., 2024). These neurons include dI4 neurons that have been suggested to inhibit neurotransmission from central terminals of nociceptive afferents to limit activation of superficial neurons primarily involved in pain-related reflexive behaviour and increase the activation of multimodal neurons downstream (Woller et al., 2017; Gatto et al., 2021; Lavaud et al., 2024). Premotor neurons that integrate multiple sensory modalities and play a role in more complex behaviours were also implicated in this form of motor adaptation by modulating motoneuron output (Gatto et al., 2021; Lavaud et al., 2024). Neurons expressing Tlx3, which includes populations of dI3 and dI5 neurons, were specifically identified as premotor neurons that mediate this function. Lastly, to facilitate the recall of this motor behaviour over longer periods, premotor inhibitory Renshaw cells were implicated for their modulation of the activity of agonist and antagonist muscles (Lavaud et al., 2024).

Given that these previous studies of motor adaptation at the spinal level mainly targeted pain pathways through stimulation of high-threshold afferents and ablation of circuits primarily processing nociceptive information (Buerger and Fennessy, 1971; Xu et al., 2008; Baumbauer et al., 2009; Lavaud et al., 2024), we asked whether dI3 neurons, a spinal population previously implicated in these behaviours (Lavaud et al., 2024) facilitate these sustained adaptive motor behaviours in response to low-threshold stimuli. If low-threshold mechanosensitive inputs could mediate motor adaptation through spinal circuits, dI3 neurons would likely be involved considering their strong inputs from proprioceptive and low-threshold mechanoreceptive afferents and the involvement of these spinal neurons in corrective motor behaviours and recovery after spinal cord injury (Bui et al., 2013, 2016; Ozyurt et al., 2025).

### Sustained motor adaptations rely on contingent sensory stimuli

We first tested whether motor adaptation in response to stimulation was specific to the animal whose toe is triggering stimulation. Similar to the approach adopted by Lavaud et al. (2024), “learner” mice were paired with “control” mice that received stimulation of the saphenous nerve that was contingent on the toe elevation of the “learner” mice. In learner mice, both the average toe position and the time above threshold were significantly higher during the trial and post-trial phases compared to pre-trial. In contrast, control mice did not show changes in these parameters across phases; therefore, these findings suggest that motor adaptation requires context-dependent and relevant stimuli. This finding is consistent with prior work demonstrating that motor adaptation in the spinal cord requires a contingent relationship between body positioning or muscle activity and stimulation (Buerger and Fennessy, 1971; Lavaud et al., 2024).

### Activation of low-threshold mechanoreceptors (LTMRs) alone is sufficient to drive spinal motor learning

While previous studies have demonstrated the capacity for sustained motor adaptations primarily through nociceptive pathways (Buerger & Fennessy, 1971; Lavaud et al., 2024), the precise control of stimulation location and intensity was often lacking, making it difficult to attribute learning to specific sensory modalities. In this study, we addressed this limitation by stimulating the saphenous nerve, a cutaneous hindlimb nerve containing Aβ, Aδ, and C fibres (Abraira & Ginty, 2013; Milenkovic et al., 2008), at defined current intensities to dissect the contributions of different afferent fibers.

We investigated whether selective activation of low-threshold fibres is sufficient to induce spinal motor learning. We determined the threshold for eliciting motor responses in our mice to be 0.3 mA, putatively targeting these low-threshold Aβ fibres that are comprised of LTMRs (Koga et al., 2005; Abraira and Ginty, 2013). Here, both toe position and the proportion of time above threshold were significantly increased during and after the trial compared to pre-trial baseline, demonstrating that the activation of LTMRs alone is sufficient to elicit these sustained motor adaptations. We also performed stimulations at higher intensities (0.6 mA and 1.2 mA) to also recruit Aδ and C fibers, which are known to play a role in these sustained motor behaviours (Buerger and Fennessy, 1971; Wall and Woolf, 1984; Joynes et al., 2004; Baumbauer et al., 2009; Lavaud et al., 2024); however, given that our low-threshold stimulations already induced reliable motor adaptations, it is difficult to differentiate their relative contribution from these high-threshold afferents in our paradigm. Future studies utilizing targeted stimulation of these distinct populations of sensory neurons in the dorsal root ganglion (DRG) using genetic tools will provide more information regarding their specific contributions to these sustained behaviours, and possible differences in downstream circuits.

### dI3s are required for motor learning in the closed-loop paradigm

To directly test the requirement for dI3 neuron activity, we used DREADD (hM4Di) receptors, which have been shown to reliably hyperpolarize targeted neurons when activated by a designer drug (Roth, 2016). This approach allowed us to reversibly silence dI3 neurons within the same animal, enabling within-subject comparisons of motor learning with and without dI3 activity. Our method provided temporal control over neuronal activity; however, because both dI3s and a subset of nociceptive sensory afferents (Aδ and C fibres) express Isl1 and Vglut2 (Goltash et al., 2025; Lagerström et al., 2010), our intersectional genetic strategy resulted in hM4Di expression in both populations, which was a potential confounding variable for high-threshold stimulations. Because Aδ and C fibres require approximately at least 2.5 and 4 times this threshold to be activated, respectively (Koga et al., 2005), our closed-loop trials at 0.3 mA stimulation intensity, which is at the behavioral threshold, allowed us to investigate the contribution of dI3s in response to low-threshold stimuli without any confounding variables.

As mentioned previously, saline administered mice had robust sustained adaptations. Specifically, both the toe position and time above threshold increased during and after the trial. In contrast, after dI3 silencing, this ability was abolished as the number of stimulations required to elevate the toe (stimulation count) was significantly higher, and the total time toe elevation was maintained above threshold was significantly shorter during and after the trial. Collectively, these results strongly support the conclusion that dI3s are required for the acquisition of sustained motor adaptations by integrating low-threshold cutaneous and likely proprioceptive input (Bui et al., 2013; Ozyurt et al., 2025).

### Model of dI3s in motor adaptation

From a control theory perspective, how might dI3s contribute to motor adaptation? Following a previous proposed framework for spinal motor learning (Brownstone et al., 2015), we propose that dI3s function as comparators, integrating feedforward and feedback signals to guide adaptive motor output. While this framework is well-established in studies of cerebellar motor learning, similar principles can be applied to the spinal cord (Brownstone et al., 2015). Brownstone et al. (2015) argued that motor adaptation at the level of spinal circuits is not limited to reflex execution but can support experience-dependent learning through internal evaluative mechanisms. Central to their model is a comparator circuit that receives both an efference copy of the motor command (feedforward signal) and actual sensory feedback. When there is a mismatch between predicted and actual feedback, a sensory prediction error is generated and used to update motor output, allowing the refinement of motor patterns over time. For dI3s to act as comparators, they must receive both sensory input from peripheral afferents and feedforward signals related to motor intent.

A recent preprint described connectivity patterns of dI3s that suggest they form a comparator module (Ozyurt et al., 2025). Indeed, dI3s receive input from proprioceptive, low-threshold cutaneous, putatively nociceptive afferents, Renshaw cells, and project to motoneurons (Bui et al., 2016; Goltash et al., 2023; Nasiri et al., 2024; Ozyurt et al., 2025). These properties position dI3s to detect mismatches between expected and actual feedback and support their proposed role in sustained motor adaptation, which is consistent with both our findings and those of Lavaud et al. (2024).

In our experiments, stimulation is delivered when muscle activity causes the toe to drop below a vertical threshold. The excitatory proprioceptive feedback to dI3s from the lowering of the limb is neutralized by the predicted motoneuron output, conveyed by Renshaw cell inhibitory efference copies to dI3s. However, stimulation of the saphenous nerve would provide cutaneous feedback to dI3s, which in turn introduces a mismatch between predicted and actual feedback. We hypothesize that dI3s detect this mismatch and drive adaptive changes in motor output to eliminate the error, thereby elevating the toe position. As adaptation progresses and the hindlimb adjusts to keep the toe above threshold, stimulation stops, decreasing excitatory drive to dI3s which aligns predicted and actual feedback; however, to maintain this equilibrium, synaptic or intrinsic mechanisms that sustain muscle activation are required—either at the level of dI3s or motoneurons.

### Proposed mechanisms by which dI3 neurons facilitate sustained motor adaptation

A recent preprint suggests that dI3 neurons form recurrent excitatory connections with motoneurons (Ozyurt et al., 2025), which could act to sustain motor activity to maintain an elevated toe position in the absence of any supraspinal or sensory excitation. This recurrent connectivity could be sufficient but might require other mechanisms to maintain motor activity for minutes. Motoneurons are well known for having persistent inward currents that can generate sustained firing activity such as plateau potentials (Schwindt and Crill, 1980; Hounsgaard and Kiehn, 1989; Bennett et al., 1998; Lee and Heckman, 2001). These motoneuron plateau potentials could be initiated by high frequency stimulation of muscle or sensory afferents (Collins et al., 2001). In addition, spinalization can facilitate the appearance of persistent inward currents in motoneurons through changes in serotonergic function in the spinal cord (Murray et al., 2010, 2011). Activation of NMDA receptors, involved with motor learning at the spinal level (Joynes et al., 2004), can also facilitate the activation of persistent inward currents (Manuel et al., 2012). Thus, motor adaptation produced by sustained muscle activation through persistent motoneuron activity could involve persistent inward currents in motoneurons, putatively activated by repetitive synaptic excitation from dI3 neurons (Bui et al., 2013; Nasiri et al., 2024). This does not preclude dI3 neurons themselves, expressing persistent inward currents. While persistent inward currents in dI3 neurons have not been reported, the presence of these currents in spinal interneurons has been shown in the intact (Theiss et al., 2007; Dai and Jordan, 2010; Singh et al., 2025) and injured spinal cord (Thaweerattanasinp et al., 2020). Therefore, dI3 neurons may play an active role in sustaining motor activity leading to relatively long-lasting motor adaptation, going beyond their role of relaying low-threshold mechanosensitive and proprioceptive inputs to motoneurons.

## ACKNOWLEDGEMENTS

We thank Sara Goltash and Julie Tremblay for excellent animal care, Aya Takeoka for discussion and helpful suggestions, and John Lewis and Simon Chen for comments on the manuscript.

## AUTHOR CONTRIBUTIONS

EUK: contributed to the conceptualization, data collection, analysis, figure preparation, and writing of the manuscript.

SN: contributed to conceptualization, analysis, figure presentation, and writing of the manuscript.

SAC and LC: contributed to analysis, figure preparation, and writing of the manuscript. AML: contributed to data collection, conceptualization, data analysis.

TVB: contributed to the conceptualization, supervision, data analysis, and writing of the manuscript.

## DATA AVAILABILITY

Data will be made available upon request.

## REFERENCES

Abraira VE, Ginty DD (2013) The sensory neurons of touch. Neuron 79:618–639.

Agashkov K, Krotov V, Krasniakova M, Shevchuk D, Andrianov Y, Zabenko Y, Safronov BV, Voitenko N, Belan P (2019) Distinct mechanisms of signal processing by lamina I spino-parabrachial neurons. Sci Rep 9:19231.

Akay T (2014) Long-term measurement of muscle denervation and locomotor behavior in individual wild-type and ALS model mice. J Neurophysiol 111:694–703.

Akay T, Acharya HJ, Fouad K, Pearson KG (2006) Behavioral and Electromyographic Characterization of Mice Lacking EphA4 Receptors. J Neurophysiol 96:642–651.

Bastian AJ, Martin TA, Keating JG, Thach WT (1996) Cerebellar ataxia: abnormal control of interaction torques across multiple joints. J Neurophysiol 76:492–509.

Baumbauer KM, Huie JR, Hughes AJ, Grau JW (2009) Timing in the absence of supraspinal input II: regularly spaced stimulation induces a lasting alteration in spinal function that depends on the NMDA receptor, BDNF release, and protein synthesis. J Neurosci Off J Soc Neurosci 29:14383–14393.

Bennett DJ, Hultborn H, Fedirchuk B, Gorassini M (1998) Synaptic activation of plateaus in hindlimb motoneurons of decerebrate cats. J Neurophysiol 80:2023–2037.

Bouyer LJ, Rossignol S (2003) Contribution of cutaneous inputs from the hindpaw to the control of locomotion. II. Spinal cats. J Neurophysiol 90:3640–3653.

Brownstone RM, Bui TV, Stifani N (2015) Spinal circuits for motor learning. Curr Opin Neurobiol 33:166–173.

Buerger AA, Fennessy A (1970) Learning of leg position in chronic spinal rats. Nature 225:751–752.

Buerger AA, Fennessy A (1971) Long-term alteration of leg position due to shock avoidance by spinal rats. Exp Neurol 30:195–211.

Bui TV, Akay T, Loubani O, Hnasko TS, Jessell TM, Brownstone RM (2013) Circuits for grasping: spinal dI3 interneurons mediate cutaneous control of motor behavior. Neuron 78:191–204.

Bui TV, Stifani N, Akay T, Brownstone RM (2016) Spinal microcircuits comprising dI3 interneurons are necessary for motor functional recovery following spinal cord transection. eLife 5:1–20.

Collins DF, Burke D, Gandevia SC (2001) Large Involuntary Forces Consistent with Plateau-Like Behavior of Human Motoneurons. J Neurosci 21:4059–4065.

Crown ED, Ferguson AR, Joynes RL, Grau JW (2002) Instrumental learning within the spinal cord: IV. Induction and retention of the behavioral deficit observed after noncontingent shock. Behav Neurosci 116:1032–1051.

Dai Y, Jordan LM (2010) Multiple patterns and components of persistent inward current with serotonergic modulation in locomotor activity-related neurons in Cfos-EGFP mice. J Neurophysiol 103:1712–1727.

Delile J, Rayon T, Melchionda M, Edwards A, Briscoe J, Sagner A (2019) Single cell transcriptomics reveals spatial and temporal dynamics of gene expression in the developing mouse spinal cord. Dev Camb Engl 146:dev173807.

Dougherty KJ, Zagoraiou L, Satoh D, Rozani I, Doobar S, Arber S, Jessell TM, Kiehn O (2013) Locomotor Rhythm Generation Linked to the Output of Spinal Shox2 Excitatory Interneurons. Neuron 80:920–933.

Duan B, Cheng L, Bourane S, Britz O, Padilla C, Garcia-Campmany L, Krashes M, Knowlton W, Velasquez T, Ren X, Ross SE, Lowell BB, Wang Y, Goulding M, Ma Q (2014) Identification of spinal circuits transmitting and gating mechanical pain. Cell 159:1417–1432.

Fillingim RB, King CD, Ribeiro-Dasilva MC, Rahim-Williams B, Riley JL (2009) Sex, Gender, and Pain: A Review of Recent Clinical and Experimental Findings. J Pain 10:447–485.

Gatto G, Bourane S, Ren X, Di Costanzo S, Fenton PK, Halder P, Seal RP, Goulding MD (2021) A Functional Topographic Map for Spinal Sensorimotor Reflexes. Neuron 109:91–104.e5.

Goltash S, Khodr R, Bui TV, Laliberte AM (2025) An optogenetic mouse model of hindlimb spasticity after spinal cord injury. Exp Neurol 386:115157.

Goltash S, Stevens SJ, Topcu E, Bui TV (2023) Changes in synaptic inputs to dI3 INs and MNs after complete transection in adult mice. Front Neural Circuits 17 Available at: https://www.frontiersin.org/articles/10.3389/fncir.2023.1176310 [Accessed August 27, 2023].

Grau JW (2014) Learning from the spinal cord: How the study of spinal cord plasticity informs our view of learning. Neurobiol Learn Mem 108:155–171.

Hounsgaard J, Kiehn O (1989) Serotonin-induced bistability of turtle motoneurones caused by a nifedipine-sensitive calcium plateau potential. J Physiol 414:265–282.

Joynes RL, Janjua K, Grau JW (2004) Instrumental learning within the spinal cord: VI. The NMDA receptor antagonist, AP5, disrupts the acquisition and maintenance of an acquired flexion response. Behav Brain Res 154:431–438.

Kendroud S, Fitzgerald LA, Murray IV, Hanna A (2025) Physiology, Nociceptive Pathways. In: StatPearls. Treasure Island (FL): StatPearls Publishing. Available at: http://www.ncbi.nlm.nih.gov/books/NBK470255/ [Accessed July 7, 2025].

Koch SC, Del Barrio MG, Dalet A, Gatto G, Günther T, Zhang J, Seidler B, Saur D, Schüle R, Goulding M (2017) RORβ Spinal Interneurons Gate Sensory Transmission during Locomotion to Secure a Fluid Walking Gait. Neuron 96:1419–1431.e5.

Koga K, Furue H, Rashid MH, Takaki A, Katafuchi T, Yoshimura M (2005) Selective activation of primary afferent fibers evaluated by sine-wave electrical stimulation. Mol Pain 1:13.

Lagerström MC, Rogoz K, Abrahamsen B, Persson E, Reinius B, Nordenankar K, Ölund C, Smith C, Mendez JA, Chen Z-F, Wood JN, Wallén-Mackenzie Å, Kullander K (2010) VGLUT2-Dependent Sensory Neurons in the TRPV1 Population Regulate Pain and Itch. Neuron 68:529–542.

Lavaud S, Bichara C, D’Andola M, Yeh S-H, Takeoka A (2024) Two inhibitory neuronal classes govern acquisition and recall of spinal sensorimotor adaptation. Science 384:194–201.

Lee RH, Heckman CJ (2001) Essential role of a fast persistent inward current in action potential initiation and control of rhythmic firing. J Neurophysiol 85:472–475.

Li L, Rutlin M, Abraira VE, Cassidy C, Kus L, Gong S, Jankowski MP, Luo W, Heintz N, Koerber HR, Woodbury CJ, Ginty DD (2011) The Functional Organization of Cutaneous Low-Threshold Mechanosensory Neurons. Cell 147:1615–1627.

Light AR, Perl ER (1979) Spinal termination of functionally identified primary afferent neurons with slowly conducting myelinated fibers. J Comp Neurol 186:133–150.

Manuel M, Li Y, ElBasiouny SM, Murray K, Griener A, Heckman CJ, Bennett DJ (2012) NMDA induces persistent inward and outward currents that cause rhythmic bursting in adult rodent motoneurons. J Neurophysiol 108:2991–2998.

Mayer WP, Akay T (2018) Stumbling corrective reaction elicited by mechanical and electrical stimulation of the saphenous nerve in walking mice. J Exp Biol 221:jeb178095.

Mogil JS (2012) Sex differences in pain and pain inhibition: multiple explanations of a controversial phenomenon. Nat Rev Neurosci 13:859–866.

Morton SM, Bastian AJ (2006) Cerebellar contributions to locomotor adaptations during splitbelt treadmill walking. J Neurosci 26:9107–9116.

Murray KC, Nakae A, Stephens MJ, Rank M, D’Amico J, Harvey PJ, Li X, Harris RL, Ballou EW, Anelli R, Heckman CJ, Mashimo T, Vavrek R, Sanelli L, Gorassini MA, Bennett DJ, Fouad K (2010) Recovery of motoneuron and locomotor function after spinal cord injury depends on constitutive activity in 5-HT2C receptors. Nat Med 16:694–700.

Murray KC, Stephens MJ, Ballou EW, Heckman CJ, Bennett DJ (2011) Motoneuron excitability and muscle spasms are regulated by 5-HT2B and 5-HT2C receptor activity. J Neurophysiol 105:731–748.

Nasiri S, Laliberte AM, Bui TV (2024) Mapping of dI3 neuron sensorimotor circuits across the cervical and lumbar spinal cord.: 2024.11.17.624039 Available at: https://www.biorxiv.org/content/10.1101/2024.11.17.624039v1 [Accessed August 24, 2025].

Nguyen-Vu TDB, Kimpo RR, Rinaldi JM, Kohli A, Zeng H, Deisseroth K, Raymond JL (2013) Cerebellar Purkinje cell activity drives motor learning. Nat Neurosci 16:1734– 1736.

Ozyurt MG, Chiasson S, Laliberte AM, Nascimento F, Khan E, Mayer WP, Bhumbra GS, Akay T, Bui TV, Beato M, Brownstone RM, Ronzano R (2025) Evidence of spinal cord comparator modules for rapid corrections of movements.: 2025.08.27.672590 Available at: https://www.biorxiv.org/content/10.1101/2025.08.27.672590v1 [Accessed September 4, 2025].

Pan Y-Z, Pan H-L (2004) Primary Afferent Stimulation Differentially Potentiates Excitatory and Inhibitory Inputs to Spinal Lamina II Outer and Inner Neurons. J Neurophysiol 91:2413–2421.

Pearson KG, Acharya H, Fouad K (2005) A new electrode configuration for recording electromyographic activity in behaving mice. J Neurosci Methods 148:36–42.

Roth BL (2016) DREADDs for Neuroscientists. Neuron 89:683–694.

Russ DE, Cross RBP, Li L, Koch SC, Matson KJE, Yadav A, Alkaslasi MR, Lee DI, Le Pichon CE, Menon V, Levine AJ (2021) A harmonized atlas of mouse spinal cord cell types and their spatial organization. Nat Commun 12:5722.

Schwindt PC, Crill WE (1980) Properties of a persistent inward current in normal and TEA-injected motoneurons. J Neurophysiol 43:1700–1724.

Singh S, Shevtsova NA, Yao L, Rybak IA, Dougherty KJ (2025) Properties of rhythmogenic currents in spinal Shox2 interneurons across postnatal development. J Physiol 603:3201–3221.

Sławińska U, Majczyński H, Dai Y, Jordan LM (2012) The upright posture improves plantar stepping and alters responses to serotonergic drugs in spinal rats. J Physiol 590:1721– 1736.

Smith MA, Shadmehr R (2005) Intact Ability to Learn Internal Models of Arm Dynamics in Huntington’s Disease But Not Cerebellar Degeneration. J Neurophysiol 93:2809– 2821.

Synofzik M, Lindner A, Thier P (2008) The Cerebellum Updates Predictions about the Visual Consequences of One’s Behavior. Curr Biol 18:814–818.

Taylor JA, Klemfuss NM, Ivry RB (2010) An Explicit Strategy Prevails When the Cerebellum Fails to Compute Movement Errors. The Cerebellum 9:580–586.

Thaweerattanasinp T, Birch D, Jiang MC, Tresch MC, Bennett DJ, Heckman CJ, Tysseling VM (2020) Bursting interneurons in the deep dorsal horn develop increased excitability and sensitivity to serotonin after chronic spinal injury. J Neurophysiol 123:1657–1670.

Theiss RD, Kuo JJ, Heckman CJ (2007) Persistent inward currents in rat ventral horn neurones. J Physiol 580:507–522.

Todd AJ (2017) Identifying functional populations among the interneurons in laminae I-III of the spinal dorsal horn. Mol Pain 13:1744806917693003.

Wall PD, Woolf CJ (1984) Muscle but not cutaneous C-afferent input produces prolonged increases in the excitability of the flexion reflex in the rat. J Physiol 356:443–458.

Woller SA, Eddinger KA, Corr M, Yaksh TL (2017) An overview of pathways encoding nociception. Clin Exp Rheumatol 35 Suppl 107:40–46.

Xu Y, Lopes C, Qian Y, Liu Y, Cheng L, Goulding M, Turner EE, Lima D, Ma Q (2008) Tlx1 and Tlx3 coordinate specification of dorsal horn pain-modulatory peptidergic neurons. J Neurosci Off J Soc Neurosci 28:4037–4046.

Zhang J, Lanuza GM, Britz O, Wang Z, Siembab VC, Zhang Y, Velasquez T, Alvarez FJ, Frank E, Goulding M (2014) V1 and V2b interneurons secure the alternating flexor-extensor motor activity mice require for limbed locomotion. Neuron 82:138–150.

Zhang Y, Narayan S, Geiman E, Lanuza GM, Velasquez T, Shanks B, Akay T, Dyck J, Pearson K, Gosgnach S, Fan CM, Goulding M (2008) V3 spinal neurons establish a robust and balanced locomotor rhythm during walking. Neuron 60:84–96.

Zimmerman AL, Kovatsis EM, Pozsgai RY, Tasnim A, Zhang Q, Ginty DD (2019) Distinct Modes of Presynaptic Inhibition of Cutaneous Afferents and Their Functions in Behavior. Neuron 102:420–434.e8.

